# Shared neural representations of cognitive conflict and negative affect in the medial frontal cortex

**DOI:** 10.1101/824839

**Authors:** Luc Vermeylen, David Wisniewski, Carlos González-García, Vincent Hoofs, Wim Notebaert, Senne Braem

## Abstract

Influential theories of medial frontal cortex (MFC) function suggest that the MFC registers cognitive conflict as an aversive signal, but no study directly tested this idea. Instead, recent studies suggested that non-overlapping regions in the MFC process conflict and affect. In this pre-registered human fMRI study, we used multivariate pattern analyses to identify which regions respond similarly to conflict and aversive signals. The results reveal that, of all conflict- and value-related regions, the ventral pre-supplementary motor area (or dorsal anterior cingulate cortex) showed a shared neural pattern response to different conflict and affect tasks. These findings challenge recent conclusions that conflict and affect are processed independently, and provide support for integrative views of MFC function.

## Introduction

The medial frontal cortex (MFC), and dorsal anterior cingulate cortex (dACC) in particular, have been implicated in various psychological processes such as cognitive control, somatic pain, emotion regulation, reward learning and decision making (Ebitz & Hayden, 2016; Heilbronner & Hayden, 2016; Shackman et al., 2011). In the domain of cognitive control, the dACC is consistently activated by cognitive conflict, that is, the simultaneous activation of mutually incompatible stimulus, task, or response representations (Botvinick et al., 2001). However, other studies also showed dACC’s involvement during the evaluation of negative outcomes, such as reductions in reward (Gehring & Willoughby, 2002), negative feedback (Nieuwenhuis et al., 2004) and pain (Rainville, 2002). Therefore, more integrative accounts of the dACC started to redescribe its role in conflict monitoring as detecting a domain-general aversive learning signal that can bias behavior away from the source of conflict (Botvinick, 2007; Shackman et al., 2011; Shenhav et al., 2013, 2016). In other words, the dACC is thought to register “cognitive” conflict as an aversive event. In accordance with this idea, recent studies have supported the presence of a behavioral bias to avoid conflict, and have also shown that humans automatically evaluate conflict as negative (Dignath et al., 2020; Dreisbach & Fischer, 2015; Inzlicht et al., 2015).

If conflict is registered as an aversive event in the dACC, one intriguing possibility is that conflict and negative affect are encoded similarly in dACC (“shared or overlapping representations”, Kragel et al., 2018). This conjecture also relates to a broader debate on the functional organization of the MFC in regard to the psychological domains of cognitive control, negative affect and pain (Inzlicht et al., 2015; Lieberman & Eisenberger, 2015; Shackman et al., 2011). Towards the end of the last century, cognitive neuroscientists argued for the functional segregation of affect (ventral part of the ACC) and cognitive control (dorsal part of the ACC) in the MFC (Bush et al., 2000; Devinsky et al., 1995). In contrast, an influential meta-analysis by Shackman and colleagues (2011) seemed to contradict this idea by showing substantial overlap in the dorsal part of the ACC for the three-way conjunction of negative affect, conflict and pain (Shackman et al., 2011). Similarly, one previous study tried to investigate the overlap in activation between cognitive conflict and negative affect by using a repetition suppression procedure, and found that dACC showed an attenuated response to negative affect following cognitive conflict (Braem et al., 2017).

In more recent years, however, other studies failed to provide evidence for such functional integration. For example, a number of recent studies and meta-analyses demonstrated that distinct rather than overlapping parts of the MFC are associated with cognitive conflict and pain processing (De La Vega et al., 2016; Jahn et al., 2016; Lieberman et al., 2016). Similarly, one recent mega-analysis study reported a multivariate pattern analysis (MVPA) on full activation maps from 18 studies to assess the similarity of patterns evoked by different domains. This study also did not observe overlap between the activation patterns evoked by cognitive control, pain, and negative emotion in the MFC (Kragel et al., 2018). Taken together, current studies seem to be at odds with integrative views of MFC (Botvinick, 2007; Brown & Alexander, 2017; Calhoun & Hayden, 2015; Heilbronner & Hayden, 2016; Shackman et al., 2011; Shenhav et al., 2013, 2016), which aim to explain the various responses of dACC by one underlying process (e.g., avoidance learning, value estimation, surprise processing). However, these studies relied on group-averaged (often univariate) activation differences originating from distinct paradigms. For example, the mega-analysis by Kragel and colleagues (2018) used a context-insensitive “cognitive control” signal (i.e., across working memory, response inhibition, and conflict processing tasks) considering paradigms from multiple studies that differ in experimental control. Moreover, many of these previous studies made use of intense pain responses that could mask similarities with the arguably subtler affective evaluation of cognitive conflict.

Here, we took a different approach and developed a more targeted and well-controlled within-subjects test of shared neural representations of conflict and negative affect. Namely, by using multivariate cross-classification analyses, we assessed whether and where a classifier algorithm trained to discern conflict (incongruent vs congruent events) can successfully predict affect (negative vs positive events), and vice versa. Successful classification (i.e., classification above chance) would be indicative of a similarity between the neural pattern response, and thus a shared representational code between these two domains (Kaplan et al., 2015; Wisniewski, 2018). To test our predictions, 38 human subjects performed a color Stroop (Stroop, 1935) and flanker task (Eriksen & Eriksen, 1974) in the conflict domain, and two closely matched tasks in the affective domain (Fig. 1A). Importantly, we used two tasks in each domain in order to first demonstrate an abstract or generalizable representation of conflict (and affect), that is independent of conflict type (and affect source) (Jiang & Egner, 2014). Conflict and affect-related brain signals were modeled using run-wise beta images retrieved from single subject (first-level) analysis and were then used to perform a five-fold leave-one-run-out cross-classification analysis using a linear Support Vector Machine (see Methods).

**Figure 1.**
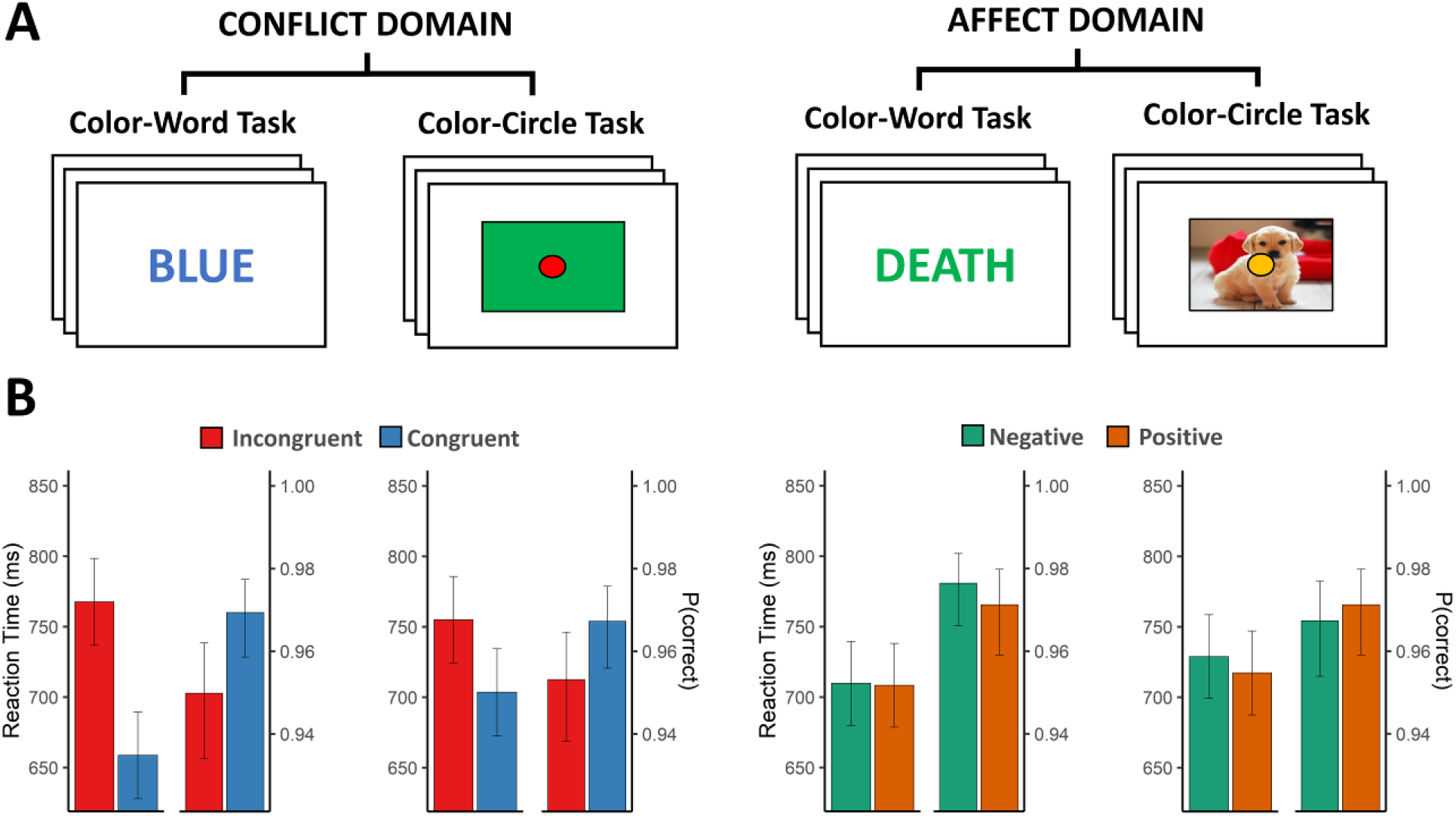
Task Design and Behavioral Data. **(A)** Task design. Subjects either judged the color of words or the color of circles. In the conflict domain, the color either matched or mismatched with word meaning or background color creating congruent or incongruent conditions, respectively. In the affective domain, positive or negative words and pictures were used to create the respective conditions. Note that regardless of the domain, subjects always had to judge the color of the word or the circle. These four task contexts were presented block-wise (with order counterbalanced) in each run. **(B)** Reaction time and accuracy for the corresponding four task contexts. Error bars denote the 95% CI.

Next, we performed preregistered Region of Interest (ROI, Fig. 2D) and whole brain searchlight analyses (Supplementary Table 1), and report accuracy-minus-chance values (chance level: 50 %) for each searchlight sphere or ROI (Amygdala, Anterior Cingulate Cortex [ACC], dACC/pre-SMA, Anterior Insula [AI], Posterior Cingulate Cortex [PCC], Ventral Striatum [VS], and the ventromedial Prefrontal Cortex [vmPFC]). To construct the ROIs, we entered the name of the anatomical area as a search term in Neurosynth (Yarkoni et al., 2011) and constructed 10mm spheres around the peak z-value of the corresponding meta-analytical activation map (see Methods and ^1^). For our main set of ROI analyses (i.e. successful cross-domain cross-task classification), we report the uncorrected p-values and Bayes Factors (always one-tailed; is accuracy-minus-chance larger than 0?), as well as the Bonferroni-corrected p-values that control for the fact that we analyzed seven ROIs (i.e., multiplying the uncorrected p-value by 7). Please note that although our literature study above focuses on the MFC, we wanted to ensure in our preregistration that other regions could be evaluated as well. For the other (secondary) set of ROI analyses, we present uncorrected p-values, but these results can be more strictly evaluated in light of the adjusted Bonferroni alpha level for seven tests (α = .00714).

**Figure 2.**
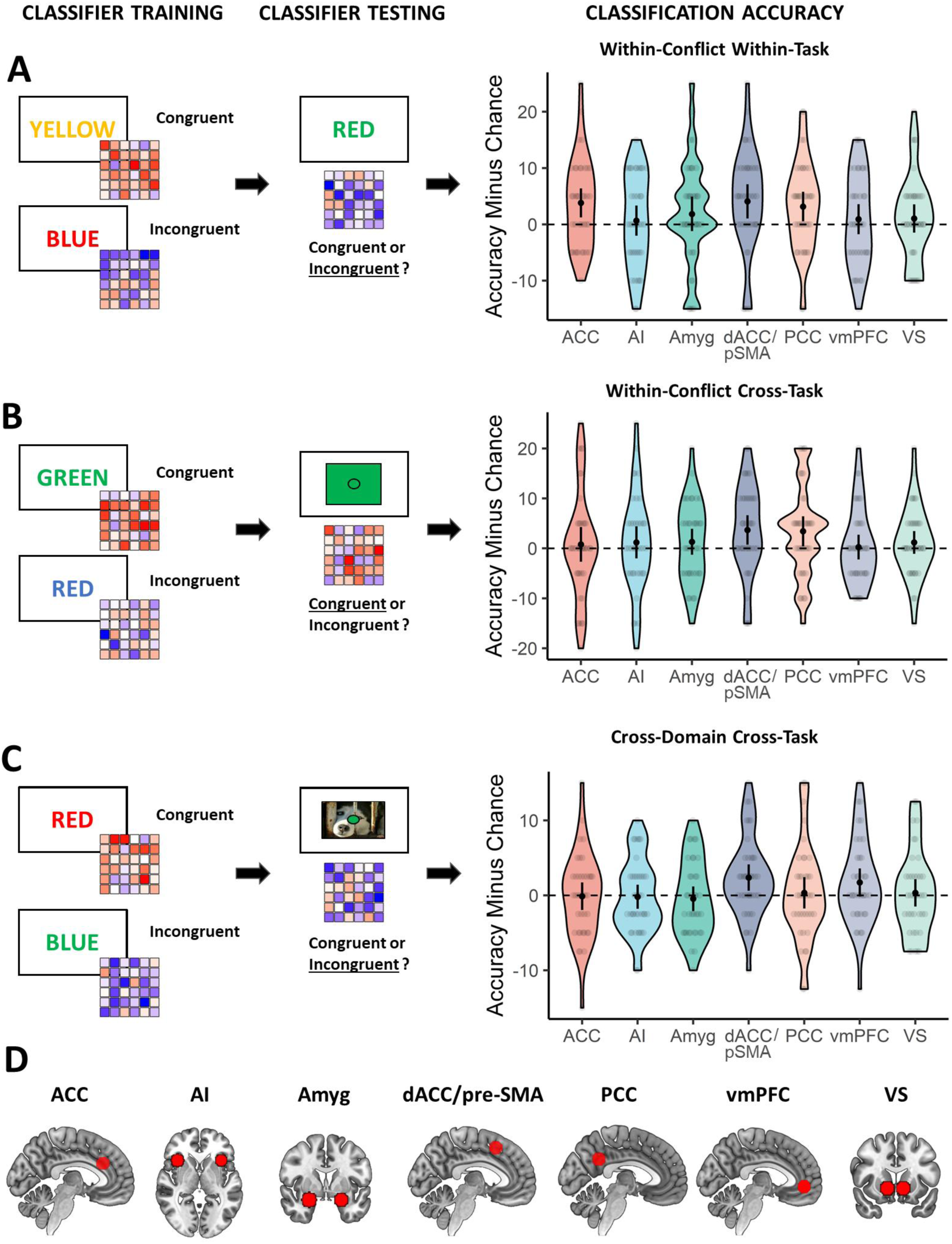
Main Results. **(A)** Training and testing the classifier within the conflict domain, within the same task. **(B)** Training the classifier on one conflict task and testing its performance on another conflict task. **(C)** Training the classifier to discern conflict and testing its performance on classifying affect in another task (and vice versa). **(D)** ROIs: Anterior Cingulate Cortex (ACC), Anterior Insula (AI), Amygdala (Amyg), dorsal Anterior Cingulate Cortex/pre-Supplementary Motor Area (dACC/pre-SMA or dACC/pSMA), Posterior Cingulate Cortex (PCC), ventromedial Prefrontal Cortex (vmPFC), Ventrial Striatum (VS). Black dots and error bars represent mean and ± 95 CI respectively; transparent dots represent individual data points; the shape of the violin shows the distribution of the data.

## Results

We first analyzed the behavioral data to check whether the conflict task showed the typical congruency effects, and subjects processed the affective stimuli in the affective task. The behavioral data from the conflict tasks (Fig. 1B, left panels and Supplementary Table 3) showed typical congruency effects in reaction time (RT, *F*_(1,37)_=149.81, *P*<.001, *BF10*>100) and accuracy (*F*_(1,37)_=11.72, *P*=.002, *BF*=52), which were larger in the color-word task (*M*=109 ms, *T*_(68)_=13.39, *P*<.001) relative to the color-circle task (*M*=51 ms, *T*_(68)_=6.31, *P*<.001) for the RT measure (interaction between task and congruency effect: *F*_(1,37)_=35.55, *P*<.001, *BF*>100). In the affective tasks, we found slower RTs with negative relative to positive background stimuli (*F*_(1,37)_=5.74, *P*=.025, *BF*=0.38), but this was only the case for the color-circle task (*M*=12 ms, *T*_(73)_=3.12, *P*=.003) and not the color-word task (*M*=1 ms, *T*_(73)_=0.37, *P*=.710) (interaction between task and valence effect: *F*_(1,37)_=4.27, *P*=.046, *BF*=0.37) (Fig. 1B, right panels and Supplementary Table 4). No effects on accuracy were found in the affective tasks (*F*s<3.12, *P*s>.085). In the conflict tasks, catch trials were used to draw attention to the conflicting nature of the stimuli. Here, subjects had to judge the irrelevant rather than the relevant dimension (see Method). We observed above-chance catch trial performance (chance level = 25 %), which did not differ between the two conflict tasks, (χ^2^_(1)_=0.10, *P*=.755, *BF*=0.12; color-circle task: 90.7 %; color-word task 89.9 %). In the affective tasks, catch trials (where subjects had to make a valence judgement instead of a color judgement) and a post-experiment incidental memory test were used to inform processing of the (task-irrelevant) affective stimuli. We observed above-chance catch trial performance (chance level = 50%), which did not differ between the two affect tasks, (χ^2^_(1)_=0.19, *P*=.664, *BF*=0.12; color-circle task: 86.4 %; color-word task 87.8 %), and successful post-experiment recognition of the affective stimuli (Supplementary Figure 5), ensuring that subjects processed the affective pictures.

In a first set of multivariate pattern analyses, we trained and tested a classifier within-task (within the Stroop or flanker task; Fig. 2A, left panels; which regions respond to conflict within tasks?), in each of our preregistered ROIs (for analysis details, see Method). Within-task ROI analyses in the conflict domain (congruent vs. incongruent) revealed evidence for above chance-level decoding in the ACC (*Wilcoxon V*=388, *uncorrected P=*.003, *BF*=15.61, Bonferroni-adjusted α = .00714), dACC/pre-SMA (*V*=327, *P=*.009, *BF*=8.48) and PCC (*V*=346, *P=*.009, *BF*=4.57) but not in in any of the other regions (all *P*>.145, *BF*<0.61) (Fig. 2A, right panel). The decoding accuracies did not differ by task (ACC: *F*_(1,37)_=0.01, *P=*.915, *BF*=0.18, dACC/preSMA: *F*_(1,37)_=0.72, *P=*.400, *BF*=0.24, PCC: *F*_(1,37)_=0.51, *P=*.476, *BF*=0.22).

A second set of multivariate pattern analyses evaluated whether we could also cross-classify conflict signals cross-task (train and test on different tasks; which regions respond similarly to conflict independent of specific task features? Fig. 2B, left panels). To our knowledge, no study has shown a voxel pattern response to conflict that is independent of conflict task in the ACC or pre-SMA (e.g., Jiang & Egner, 2014). Here, our within-conflict cross-task ROI analyses revealed above chance level conflict decoding across tasks in the dACC/pre-SMA (*V*=283, *P=*.012, *BF*=5.57) and PCC (*V*=328, *P=*.023, *BF*=3.67) (Fig. 2B, right panel). Decoding accuracy did not differ between cross-task combination for the dACC/pre-SMA (*F*_(1,37)_=0.89, *P=*.352, *BF*=0.26), but was larger in the Stroop to flanker decoding relative to the flanker to Stroop decoding for the PCC (*F* _(1,37)_=4.35, *P=*.043, *BF*=1.21). These results were also replicated in an overall decoding approach where the classifier was trained and tested in the whole domain regardless of task (resulting in more samples to train the classifier; Supplementary Fig. 1A). Within the affective domain (positive vs. negative), we also performed similar within- and cross-task decoding analyses. However, while these analyses showed evidence for affect information in the AI and vmPFC, they did not show evidence for decoding in the ACC, dACC/pre-SMA or PCC (Supplementary Fig. 2; but see follow-up analyses below).

Finally, we investigated our main hypothesis by training a classifier on discerning conflict (incongruent vs congruent) and testing its performance on discerning affect (negative vs positive), and vice versa. For this main set of analyses, we report the uncorrected and Bonferonni-corrected p-values, and focused on the cross-domain cross-task decoding (train and test in different domains on different tasks) as this analysis controls for low-level features shared between the two tasks (Fig. 2C, left panel). The cross-domain cross-task ROI decoding revealed evidence for cross-classification in the dACC/pre-SMA (*V*=330, *P*=.007, *BF*=8.43; Fig. 2C, right panel) and vmPFC (*V*=277, *P*=.045, *BF*=1.56), which did not differ by cross-task combination (dACC/pre-SMA: *F*_(1,37)_=0.36, *P=*.551, *BF*=0.20, vmPFC: *F*_(1,37)_=0.02, *P=*.880, *BF*=0.18). Bonferroni corrected p-values controlling for the use of seven ROIs showed this result was still significant for the dACC/pre-SMA (*P*_*corr*_=.049), but not for the vmPFC (*P*_*corr*_=.316). None of the other ROIs reached significance (all *P*s>.444, *BF*s <0.24). Using the overall decoding approach (training on both tasks in one domain and testing on both tasks in the other domain), we were only able to replicate successful cross-domain decoding in dACC/pre-SMA (*V*=449, *P*=.021, *BF*=4.65; Supplementary Fig. 1C).

In what follows, we report three additional sets of control analyses. A first set was designed to evaluate the extent to which our analysis choices influenced our findings (Botvinik-Nezer et al., 2020). A second set was geared towards ruling out alternative hypotheses. Finally, a third set further zoomed in on the above observation that we could not decode affect within- or across-tasks in the dACC/pre-SMA, but did observe cross-domain decoding.

First, a number of control analyses using different analysis choices further confirmed our main finding. We replicated our results using permutation testing (see Methods), as classification rates might not match theoretical chance levels (Jamalabadi et al., 2016). This approach revealed similar results (dACC/pre-SMA: *P*=.006, *P*_*corr*_=.041, vmPFC: *P*=.034, *P*_*corr*_=.238) and further demonstrated that chance level was not inflated as the mean of the group null distribution of the accuracy minus chance measure did not differ from zero for each of the two ROIs (dACC/pre-SMA: *P*=.753; vmPFC: *P=*.267). We also replicated our main finding using different smoothing parameters (Supplementary Fig. 3), or when using Harvard-Oxford atlas ROIs (Supplementary Fig. 4) instead of ROIs retrieved from NeuroSynth. Moreover, when using a set of functionally (rather than anatomically) defined conflict-sensitive ROIs based on a recent meta-analysis (from Chen et al., 2018, see their Table 2 for MNI coordinates), we again observed evidence for cross-domain cross-task classification in the dACC/pre-SMA (*V*=450, *P*=.013, *BF*=3.75) but not for other conflict-sensitive ROIs (left MOG, right AI, left AI, left IFG, left IPL, right IPL, left MFG), except for the left AI (*V*=425, *P*=.005, *BF*=8.61). The result again replicated when using the overall decoding approach in the dACC/pre-SMA (*V*=449, *p*=.001, *BF*=41.06), but not in the left AI (*V*=335, *P=*.260, *BF*=0.34) (Supplementary Fig. 1, panel D).

Second, to further study whether the main effect was specific to the congruency identity and valence of our experimental conditions, we also tested whether task difficulty differences or differences in arousal could not explain our results. Notably, there was a small performance difference in task performance on negative versus positive affect trials. Therefore, it is possible that our cross-decoding result reflects the decoding of reaction time, rather than a shared conflict and affect signal. However, the RT effects for the affective domain were rather small (*BF*<1) and only present for one of the two affective tasks (also, note that there were no effects on accuracy). Importantly, as reported above, the cross-domain decoding accuracy did not depend on which task was used for testing. Moreover, cross-domain decoding accuracy was not higher when splitting the sample for the subsample that did show the RT effect (RT effect>0, N=28, *M*=1.96, *P*=.050, *BF*=1.66) relative to a subsample that did not show the RT effect (RT effect<0, N=10, *M* =3.50, *P*=.017, *BF*=4.58) (*F*_(1,36)_=0.59, *P=*.444), nor did the RT effect correlate with cross-domain decoding accuracy (*R*=-0.23, *P*=.162). Nevertheless, in order to control for any potential confounding effects of RT, we also ran another control analysis regressing out RT-related effects from the neural data (via parametric modulation). This analysis led to the same conclusions as above (cross-domain cross-task dACC/pre-SMA: *V*=380, *p*=.036, *BF*=1.91, overall cross-domain dACC/pre-SMA: *V*=348, *p*=.025, *BF*=2.61). Note that this does make the effect slightly smaller, which should not be surprising as this procedure also removes variance of interest (as the effectiveness of the congruency manipulation is often defined by RT differences).

Next, we also found that the cross-domain decoding was unlikely to be due to arousal. First, we were only able to decode arousal (high vs. low arousal; based on a median-split of arousal ratings, orthogonal to valence) within-tasks in the ACC (V=260, *P*=.039, *BF*=1.43) and vmPFC (V=240, *P*=.048, *BF*=1.38), but not in the dACC/pre-SMA (V=230, *P*=.082, *BF*=0.83) or other ROIs (*P*>.217, *BF*<0.39). Second, training a classifier on conflict and testing on arousal (and vice versa) did not show any evidence for cross-decoding in the dACC/pre-SMA (V=294, *P*=.404, *BF*=0.26), nor any of the other ROIs (*P*s>.139, *BF*<0.51), except for the vmPFC (*V*=401, *P*=.015, *BF*=3.25).

Third, and finally, we wanted to follow up on the unexpected result that we did not observe within and cross-task decoding of affect in the dACC/pre-SMA. One potential explanation is that the SNR was lower for our affect differences than the congruency differences. In line with this, while the univariate affect contrast (negative > positive) showed no cluster-corrected activation in the MFC (Supplementary Table 2), a large cluster of dmPFC activity does become visible, when we lower the threshold (p < .005, uncorrected). Similarly, when restricting the whole-brain decoding analysis to the MFC using the same mask as Kragel and colleagues (2018), within-affect decoding did reveal a large cluster in the MFC, overlapping with our main dACC/pre-SMA ROI, as well as the within-conflict decoding and cross-domain decoding results (see Supplementary Fig. 6). A second reason for observing weaker affect decoding signals, could be because the affect signals weakened or habituated throughout the experiment, as the same affective words and pictures were repeated in each run. Therefore, we also investigated whether affect might have been easier to decode in the beginning of the experiment, by studying cross-task affect for the first half (run 1 and 2) and second half (run 4 and 5), separately. Indeed, there was significant cross-task affect decoding in main dACC/pre-SMA ROI in the first half of the experiment (run 1 and 2, *V*=237, *P*=.019, *BF*=2.38), but not in the second half of the experiment (run 4 and 5, *V*=138, *P*=.646, *BF*=0.10). Finally, it is worth noting that our ROIs were primarily selected to detect conflict-based differences (see also our literature-based ROIs on conflict decoding, Supplementary Fig. 1, panel D), which again potentially lowered our chances to observe affect decoding. When using ROIs based on an affect processing meta-analysis (from Lindquist et al., 2016, their Table 2, cf. our conflict-sensitive ROIs), a pre-SMA ROI close to our original dACC/pre-SMA ROI (but a bit more dorsal) did show significant affect (*V*=314, *P*=.001, *BF*=39.70), conflict (*V*=308, *P*=.002, *BF*=27.19) and cross-domain decoding (*V*=375, *P*=.042, *BF*=1.52). Together, these results do suggest that there was affect decoding in, or at least around our main dACC/pre-SMA ROI.

## Discussion

Together, our results reveal that the dACC/pre-SMA shows a similar voxel pattern response to conflict and negative affect. Moreover, to the best of our knowledge, our study is also the first to show decoding of conflict across conflict tasks in the dACC/pre-SMA, suggesting a shared component in the detection of conflict across the Stroop and flanker task (Jiang & Egner, 2014).

These findings fit with integrative accounts of MFC function which propose that multiple domains (e.g., cognitive conflict, negative emotion and pain) activate a single underlying process in the MFC (e.g., adaptive control, avoidance learning, cost-benefit value estimation), and thus predict similar activation patterns across domains (Botvinick, 2007; Brown & Alexander, 2017; Calhoun & Hayden, 2015; Heilbronner & Hayden, 2016; Shackman et al., 2011; Shenhav et al., 2013, 2016). Since our study focused on the domains of cognitive conflict and negative affect, our finding is especially relevant for theories that have tried to explain dACC’s sensitivity to conflict and negative outcomes by a single unifying process (Botvinick, 2007; Shenhav 2013, 2016). These theories have proposed the idea that conflict is in itself an aversive outcome (Botvinick, 2007; Shackman et al., 2011; Shenhav et al., 2013), and the main mechanism of the dACC is to bias behavior away from any costly, demanding or suboptimal outcome (“domain-general avoidance learning”, Botvinick, 2007) or to allocate control based on the expected benefits discounted by the expected costs of controlling behavior (“cost-benefit value estimation” Shenhav et al., 2013).

Importantly, our study was not set up to disentangle the multiple candidate processes as to why conflict and negative affect would elicit a shared neural pattern response in the MFC, so while our main finding (i.e., similar voxel pattern response to conflict and negative affect) follows naturally from above-mentioned models that predict similar patterns of brain activity for different domains, it does not favor a specific mechanism (e.g. domain-general avoidance learning). Still, our finding does lend credence to the idea that that cognitive control can be understood as an emotional process (Inzlicht et al., 2015), and that conflict can be registered as an aversive event (i.e., Botvinick, 2007; Dignath et al., 2020; Dreisbach & Fischer, 2015), at least in the dACC/pre-SMA. Similarly, additional analyses further show that our effect is unlikely to be driven by general differences in reaction time or task difficulty, or differences in arousal.

Other recent studies failed to find similarities, and suggest that theories of MFC function should not look for a unitary neural implementation, but rather focus on a unified computational mechanism (Kragel et al., 2018). Our study does not argue against this idea, as shared neural “representations” in fMRI could still be driven by different local neighboring neural populations. However, we believe our findings do license consideration of a unitary neural implementation, and offer support for the idea of a unified computational mechanism. Still, one might wonder why we did, and others did not, find shared neural representations of conflict and affect. First, we employed a multivariate approach rather than a univariate approach which is more sensitive and more informative to assess similarity in brain activation (without necessarily being orthogonal to a univariate approach; Davis et al., 2014). Second, our design was specifically set up to evaluate similarity between two specific domains (conflict and negative affect) rather than very general domains (cognitive control, negative emotion, pain). Third, related to this, we used a within-subjects design (rather than the between-subject nature of meta- or mega-analyses), which allowed for more sensitive analyses and to make the different task contexts as similar as possible (high level of experimental control).

One unexpected finding relates to the fact that we did not observe a similar (significant) above-chance decoding of affect in the dACC/pre-SMA, but did observe cross-domain decoding. Although this finding was surprising to us at first, it most likely suggests differences in signal to noise ratio (SNR) between the two domains and does not invalidate the cross-domain decoding result (for a review and discussion of similar findings, see van den Hurk & de Beeck, 2019). This idea was also further corroborated by follow-up analyses which showed affect decoding in the dACC/pre-SMA when using other analysis techniques, or focusing on the first half of the experiment only. Moreover, a lower SNR in the affect domain can also be explained by the fact that affect was not relevant for the main task (which allowed us to keep the affective tasks as similar as possible to the conflict tasks).

Finally, while our analyses were based on theories and a NeuroSynth ROI of the dACC, investigating the (uncorrected) whole-brain searchlight decoding maps did reveal that the conflict, affect and cross-domain decoding all trigger a dorsomedial frontal area, that might be more accurately referred to as pre-SMA rather than dACC. This area roughly corresponds to our main dACC/pre-SMA ROI which was retrieved by searching “dACC” in NeuroSynth (Yarkoni et al., 2011), and therefore, we decided to refer to this ROI as dACC/pre-SMA. Nevertheless, this illustrates the consequent mislabeling of the dACC in the literature and we are not the first to report pre-SMA activation for domains such as cognitive control, pain and negative emotion (Brown & Alexander, 2017; De La Vega et al., 2016; Jahn et al., 2016; Kolling et al., 2016; Kragel et al., 2018; Lieberman & Eisenberger, 2015; Lindquist et al., 2016).

In sum, by using a well-controlled within-subject design and multivariate analysis techniques, we show that conflict and negative affect evoke a similar voxel pattern response in the MFC. Our finding brings important nuance to other recent studies suggesting different neural activations or representations, and helps to further constrain theories of integrative MFC function.

## Materials and Methods

### Participants

The study was pre-registered with the pre-registration template from AsPredicted.org on the Open Science Framework (https://osf.io/p5frq/). As pre-registered, 40 participants participated in our study. Two participants were excluded (one due to excessive head motion [>2.5mm translation] and one aborted the scanning session). The average age of the remaining 38 participants (13 male) was 23.71 years (*SD=*3.53, min=18, max=33). Thirty-six participants were right-handed, one was left-handed and one was ambidextrous (as assessed by the Edinburgh Handedness Inventory (Oldfield, 1971)). Every participant had normal or corrected to normal vision and reported no current or history of neurological, psychiatric or major medical disorder. Every participant gave their informed written consent before the experiment, and was paid 35 euros for participating afterwards. The study was approved by the local ethics committee (University Hospital Ghent University, Belgium).

### Experimental Paradigm

The experiment was implemented using Psychopy 2 version 1.85.2 (Peirce, 2007). On each trial, participants had to judge the color of a target stimulus in the center of the screen, using two MR-compatible response boxes (each box had two buttons) to indicate one out of four possible response options (red, blue, green and yellow). The key-to-color mapping was counterbalanced between participants. The exact features of the target stimulus varied block-wise, depending on one of four different task-contexts. Specifically, participants either had to respond to the color of words (“color-word naming task”) or respond to the color of circles (“color-circle naming task”), which both had a conflict and affective version.

The conflict-version of the color-word naming task was a Stroop task (Stroop, 1935), where the meaning of the words could either be congruent or incongruent with the actual color of the word. For example, participants could see the words “BLUE”, “RED”, “GREEN” or “YELLOW” (Dutch: “ROOD”, “BLAUW”, “GROEN” or “GEEL”) presented in a blue, red, green or yellow font. The conflict version of the color-circle naming task was essentially a color-based variant on the Eriksen flanker task (Eriksen & Eriksen, 1974), where the irrelevant feature consisted of a colored background square which could either be congruent or incongruent with the color of the circle. Here, participants could see blue, red, green or yellow circles presented on a blue, red, green or yellow background square. In both tasks, half of the trials were congruent (e.g., “RED” in a red font; a red circle presented on a red square background) while the other half of the trials were incongruent (e.g., “RED” in a blue font; a red circle on a blue square background).

The affect-versions of the color-word naming and color-circle naming tasks made use of irrelevant affective words or pictures, respectively. In the color-word naming task, 16 positive and 16 negative words were presented (Moors et al., 2013) that were matched on arousal, power, age of acquisition, Dutch word frequency (Keuleers et al., 2010), word length and grammatical category (Noun, Adjective and Verbs). The affective picture distractors in the background of the color-circle naming task were retrieved from the OASIS database (Kurdi et al., 2017). Sixteen positive and 16 negative pictures were presented that were matched on semantic category (Animals, Objects, People, Scenery) and arousal. This resulted in a total of eight conditions: congruent, incongruent, positive or negative trials, that either involved words or pictures/colored backgrounds.

Each trial started with a fixation sign (“+”) that was presented for 3 to 6.5 seconds (in steps of 0.5 s; *M=*3.5 s; drawn from an exponential distribution). Next, the target stimulus was presented for 1.5 seconds (fixed presentation time regardless of RT). In order to increase the saliency of the irrelevant dimension (conflict and affect), the onset of the affective word or picture preceded the presentation of the target feature by 200 ms during which the color of the target feature (word or circle) was white.

Participants performed five scanning runs and during each run the subjects performed each of the four task contexts in separate blocks. The order of the four blocked task contexts was fixed within participant but counterbalanced between participants. Each block hosted 32 trials (16 congruent/positive and 16 incongruent/negative) which were presented in a pseudo-random fashion with the following restriction: neither relevant nor irrelevant features of the target stimulus could be repeated. This restriction was used to investigate confound-free congruency sequence effects (see Braem et al., 2019; Schmidt, 2019; but this was not the aim of the current study and will not be discussed further). In total, each participant made 640 trials (i.e., five runs of four blocks of 32 trials).

In each task context (block), we also included one catch trial (at random, but not in the first two or last two trials of each block). In these catch trials, the presentation of the task-irrelevant word, picture, or colored square would not be followed by the presentation of the target color, and remain on screen for three seconds. Participants were instructed that during these catch trials, when no color information was present in the relevant dimension, their goal was to judge the irrelevant dimension depending on the cognitive domain. In the conflict domain, participants had to respond to the meaning of the word (“RED”, “BLUE”, “GREEN” or “YELLOW”) or to the color of the background square (red, blue, green or yellow) by using the respective key that would be used to judge the relevant dimension. In the affective domain, participants had to judge the affective word or background picture as either positive or negative by pressing all keys once or twice (response mapping for positive and negative stimuli counterbalanced between participants). The purpose of these catch trials was to increase the saliency of the irrelevant dimension.

Before the scanning session, participants were welcomed and instructed to read the informed consent after which they started practicing the experimental paradigm. After the scanning sessions, participants performed an unannounced recognition memory test on old and new affective words and pictures. Here, participants had to indicate whether they had previously seen the word or picture in the experiment (old/new judgement). The new words were matched with the old words in terms of valence, arousal, power, age of acquisition, word length, frequency, grammatical category. The new pictures were matched on valence, arousal and semantic category. In both a behavioral (n = 20) and fMRI pilot (n = 20), we already established that participants showed adequate performance on both the main task and the recognition memory task. Finally, participants completed four questionnaires (Need for Cognition, Behavioral Inhibition/Activation Scale, Positive and Negative Affect Schedule, Barret Impulsivity Scale) and were thanked for their participation. No significant correlations between these questionnaire scales and cross-classification accuracies were found (see Supplementary Table 5 and 6).

### Behavioral Data Analysis

Behavioral analyses were performed in R (RStudio version 1.1.463, www.rstudio.com). For the reaction time (RT) analyses, we removed incorrect, premature (< 150 ms), and extreme responses (RTs outside 3 SD from each condition mean for each participant). This resulted in an average of 94.42 % of the trials left for the RT analyses (*SD=*3.18, min=84.22, max=98.28). We conducted a repeated measures ANOVA on the reaction time and accuracy measure with the within-subject factors Condition (conflict domain: congruent vs. incongruent, affective domain: positive vs. negative) and Task (color-word naming vs. color-circle naming). We also assessed post-scanning recognition memory of affective stimuli with a probit generalized linear mixed effects model on the probability to say that the stimulus was ‘old’ with fixed effects for Experience (old vs. new), Valence (positive vs. negative) and Task Type (word vs. picture) and crossed random effects for Participant and Item. We also pre-registered some exclusion criteria based on behavioral performance. Participants with a mean RT outside 3 SD from the sample mean or a hit rate below 3 SD or 60 % (chance level=25 %) from the sample mean were excluded. Participants that performed poorly on the post-scanning recognition memory test, i.e., hit rate or false alarm rate outside 3 SD of the sample mean were also excluded. In the end, no exclusions based on task performance had to be made. While performance on catch trials was not a pre-registered exclusion criterion, we found that two participants responded on chance level in the catch trials of the affective domain (chance level=50 %, positive vs. negative judgement). Excluding these participants did not change our conclusions.

### fMRI data acquisition

fMRI data was collected using a 3T Magnetom Prisma MRI scanner system (Siemens Medical Systems, Erlangen, Germany), with a sixty-four-channel radio-frequency head coil. A 3D high-resolution anatomical image of the whole brain was acquired for co-registration and normalization of the functional images, using a T1-weighted MPRAGE sequence (TR=2250 ms, TE=4.18 ms, TI=900 ms, acquisition matrix=256 × 256, FOV=256 mm, flip angle=9°, voxel size=1 × 1 × 1 mm). Furthermore, a field map was acquired for each participant, in order to correct for magnetic field inhomogeneities (TR=520 ms, TE1=4.92 ms, TE2=7.38 ms, image matrix=70 x 70, FOV=210 mm, flip angle=60°, slice thickness=3 mm, voxel size=3 x 3 x 2.5 mm, distance factor=0%, 50 slices). Whole brain functional images were collected using a T2*-weighted EPI sequence (TR=1730 ms, TE=30 ms, image matrix=84 × 84, FOV=210 mm, flip angle=66°, slice thickness=2.5 mm, voxel size=2.5 x 2.5 x 2.5 mm, distance factor=0%, 50 slices) with slice acceleration factor 2 (Simultaneous Multi-Slice acquisition). Slices were orientated along the AC-PC line for each subject.

### fMRI data analysis

fMRI data analysis was performed using Matlab (version R2016b 9.1.0, MathWorks) and SPM12 (www.fil.ion.ucl.ac.uk/spm/software/spm12/). Raw data was imported according to BIDS standards (http://bids.neuroimaging.io/) and functional data was subsequently realigned, slice-time corrected, normalized (resampled voxel size 2 mm^3^) and smoothed (full-width at half maximum of 8 mm). The preprocessed data was then entered into a first-level general linear model analysis (GLM), and subsequently into a multivariate pattern analysis (MVPA(Cox & Savoy, 2003; Haxby, 2012; Haynes, 2015; Kriegeskorte et al., 2006). Results were analyzed using a mass-univariate approach. Although we pre-registered that we would not normalize and smooth the data for our classification analyses, we found that Signal-to-Noise Ratio (SNR) was significantly improved with these additional preprocessing steps (Supplementary Fig. 3A). In addition, an independent classification analysis (classifying left vs. right responses in primary motor cortex) showed that decoding accuracies were significantly higher with these additional preprocessing steps (Supplementary Fig. 3B). Knowing that decoding information in the PFC is notoriously difficult as decoding accuracies are close to chance (relative to decoding in occipitotemporal cortex (Bhandari et al., 2018), and the finding that smoothing can and does often improve SNR and decoding performance (Hendriks et al., 2017; Kamitani & Sawahata, 2010; Op de Beeck, 2010), we decided to optimize our MVPA analyses by decoding on normalized and smoothed data. For completeness, however, we also depict the results from our main cross-classification analysis for different levels of smoothing (FWHM 0, 4 and 8 mm; see Supplementary Fig. 3C).

First-level GLM analyses consisted of 5 identically modeled sessions (i.e., the five runs). Each session consists of eight regressors of interest (for the eight conditions, see above), four block regressors (to account for the blocked presentation of each combination of word versus picture versions of the conflict versus affect tasks), two nuisance regressors (that model performance errors and catch trials) and six movement regressors. The regressors were convolved with the canonical HRF. The modeled duration of the regressors of interest (the eight conditions) and nuisiance regressors (errors, catch trials) was zero, while the modeled duration of the block regressors was equal to the length of the blocks.

Next, the beta images from the first-level GLM were submitted to leave-one-run-out decoding scheme with ‘The Decoding Toolbox’ (Hebart et al., 2015) using a linear support-vector classification algorithm (C=1). We performed whole-brain searchlight decoding (sphere radius: 3 voxels; Supplementary Table 1) as well as ROI decoding (see below for ROI methods). Cross-validation decoding was conducted within the affective (positive vs. negative) and conflict (congruent vs. incongruent) domain for each task separately (“within-domain within-task classification”). To assess the generalizability of the classifier within the domain, we also conducted cross-classification analyses where we trained the classifier on one task and tested its performance on the other task for each task type combination (from color-circle naming to color-word naming and vice versa) separately (“within-domain cross-task classification”). To investigate the generalizability of these classifiers across the domain (our main hypothesis), we trained the classifier in the conflict domain and tested its performance in the affective domain, and vice versa. We conducted these analyses cross task type combinations (i.e., from color-circle naming to color-word naming, or from color-word naming to color-circle naming) to further control for low-level task features, following the same reasoning as the within-domain cross-task classification analyses (“cross-domain cross-task classification”). For each of these three decoding analyses, we also ran ANOVAs to evaluate whether the result differed depending on the task (e.g., color-circle naming versus color-word naming) or task-to-task direction (i.e., from color-circle naming to color-word naming, or from color-word naming to color-circle naming). Finally, we also report an “overall decoding” analysis, where the classifier was trained across the two task types at once, thereby ignoring whether the event featured words or pictures/colored backgrounds.

Each classification analysis resulted in ‘accuracy-minus-chance’ decoding maps for each subject. These maps were then entered into a group second-level GLM analysis in SPM12. Here, a one-sample t-test determined which voxels show significant accuracy above chance level.

Importantly, counterbalancing schemes that are not problematic for traditional analyses can introduce problems for the assessment of chance performance in MVPA (Gorgen et al., 2018). This can be the case when the counterbalancing is “broken” by splitting the runs in training and test sets given that the counterbalancing and cross-validation are situated on the same level (e.g. the within-subject level of the runs, Gorgen et al., Neuroimage 2018). However, in our study, the order of the four tasks within a scanning run was counterbalanced between participants but fixed within participant (which is where the leave one run out cross-validation occurs). Therefore, counterbalancing conditions were always matched between training and testing runs. In addition, we modeled the blocked structure of the scanning runs by adding block regressors in the first level GLM’s (before entering the beta images to the classification analyses). Therefore, it is unlikely that our counterbalancing could have introduced problems for the assessment of chance level. Nonetheless, for our main findings, this was verified by inspecting the null distributions obtained via permutation testing. Further, we did not remove univariate difference (more specifically, response differences of the same sign), as is sometimes done. Univariate differences are often a useful source of information and should not necessarily be viewed as irrelevant for multivariate analyses (Hebart & Baker, 2018). Further, we do not intend to make a special claim about fine grained subtle patterns that go beyond univariate response differences.

Next to MVPA, we also conducted classic univariate analyses. Here, we constructed a set of contrasts subtracting (A) positive from negative conditions and (B) congruent from incongruent conditions for (1) each task separately as well as across both tasks. These contrast images were then entered into a second-level analysis in which a one-sample t-test determined which voxels show significant activation for each contrast. We applied a statistical threshold of *p* < 0.001 (uncorrected) at the voxel level, and *p* < 0.05 (family-wise error corrected) at the cluster level on all analyses (Supplementary Table 2).

### ROI analyses

As part of our pre-registered main analysis plan, we conducted ROI decoding analyses. We set out to study the Amygdala, Anterior Cingulate Cortex (ACC), dorsal Anterior Cingulate Cortex/pre-SMA (dACC/pre-SMA), Anterior Insula (AI), Parietal Cingulate Cortex (PCC), Ventral Striatum (VS), and the ventromedial PFC (vmPFC). Our preregistration noted that all ROIs would be obtained from the Harvard-Oxford cortical and subcortical structural atlases, thresholded at 25%. However, as the dACC ROI was not defined in the Harvard-Oxford atlas, we decided to retrieve this ROI from Neurosynth (Yarkoni et al., 2011) by entering “dacc” as search term (returning 162 studies reporting 4547 activations). Although this ROI was based on the “dacc” search term, the peak effect of studies reporting dACC activity actually lies more dorsally than the cingulate gyrus, overlapping with the pre-SMA (Lieberman & Eisenberger, 2015). Therefore, we refer to this ROI as the dACC/pre-SMA. Next, we built a 10 mm sphere around the peak activation point in this activation map (association map). Because the dACC ROI was spherical (in contrast to the other six atlas ROIs), we decided to retrieve all ROIs from NeuroSynth for comparability. However, because our preregistration mentioned that the ROIs would be obtained from the Harvard-Oxford cortical and subcortical structural atlases, we also report the analyses where all non-dACC regions were evaluated using these ROIs, which returned similar results and did not change our main conclusions (see Supplementary Figure 4).

In addition to the pre-registered ROI analyses which were based on anatomically determined ROIs, we also ran another set of ROI analyses with functionally informed ROIs. Namely, we created 10 mm sphere ROIs for all conflict-sensitive regions based on the most recent and inclusive meta-analysis we could find on cognitive conflict (Chen et al., 2018, Table 2).

Each ROI decoding analysis returned one accuracy-minus-chance value per ROI and participant. We tested whether these values were significantly higher than zero (one-tailed) with the non-parametric Wilcoxon signed-rank test and a Bayesian t-test (using the default priors from the BayesFactor package in R; Cauchy prior width: *r=*.707). We report the Bayes Factor (BF) that quantifies the evidence for the alternative hypothesis (i.e., decoding accuracy is higher than zero). For our main hypothesis (cross-domain cross-task classification) we also report permutation tests. Here, the labels of the classification categories were shuffled (within-run only) and classification was performed for all possible permutations leading to first-level null distributions (i.e., for each subject) of accuracy-minus-chance values. A second-level null distribution was retrieved by calculating 10000 group averages (where the value for each subject is a random draw from its respective null distribution). We retrieved a p-value by dividing the number of samples above the empirical accuracy-minus-chance level by the total number of samples in the null distribution. The procedure of obtaining the second-level null distribution and p-value was repeated 1000 times after which we averaged the resulting p-values. This lead to stable p-values that are not influenced by stochasticity in the procedure. We report uncorrected as well as Bonferroni-corrected p-values for our main test. The secondary tests are not corrected as they are not the main aim of the study (but can be more strictly evaluated by handling the Bonferonni-corrected alpha: *p*<.00714). Finally, we investigated whether the significant cross-task cross-domain classification accuracy correlated with the following behavioral indices: post-scanning affective recognition memory (d-prime), congruency sequence effects in reaction time and error rate and congruency sequence effects in reaction time and error rates (p-values of reported correlations are Holm-corrected for five tests) (see Supplementary Figure 5).

## Acknowledgements

We would like to thank Tobias Egner for valuable comments on a previous draft of the manuscript. W.N., S.B. (G.0660.17N) and L.V. (11H5619N) were supported by the FWO – Research Foundation Flanders. C.G.G. was supported by the Special Research Fund of Ghent University (BOF.GOA.2017.0002.03). D.W. was supported by the FWO (FWO.KAN.2019.0023.01), and the European Union’s Horizon 2020 research and innovation program under the Marie Skłodowska-Curie grant agreement No 665501. All procedures applied in the present experiment were carried out with adequate understanding and written consent of the subjects and are in accordance with the Declaration of Helsinki.

## Author Contributions

S.B. and W.N. developed the study concept. S.B., W.N. and L.V. contributed to the study design. Data collection was performed by L.V. and V.H.. Data analysis was performed by L.V. under the supervision of S.B., D.W. and C.G.C.. The manuscript was drafted by L.V. in cooperation with S.B., W.N., D.W. and C.G.C.. All authors approved the final version of the manuscript for submission.

## Footnotes

1 We actually pre-registered that the ROIs would come from the Harvard-Oxford cortical and subcortical atlases, thresholded at 25%. However, at the time of pre-registration we did not realize that the dACC ROI has no clear analogue in the Harvard-Oxford or most other atlases. We reasoned that the most neutral way to deviate from the pre-registration was to construct spheres from peak-values of the Neurosynth maps for each respective ROI.

