## Supplementary Materials for "Shared neural representations of cognitive conflict and negative affect in the medial frontal cortex"

***Supplementary Figure 1.*** **Overall Decoding**

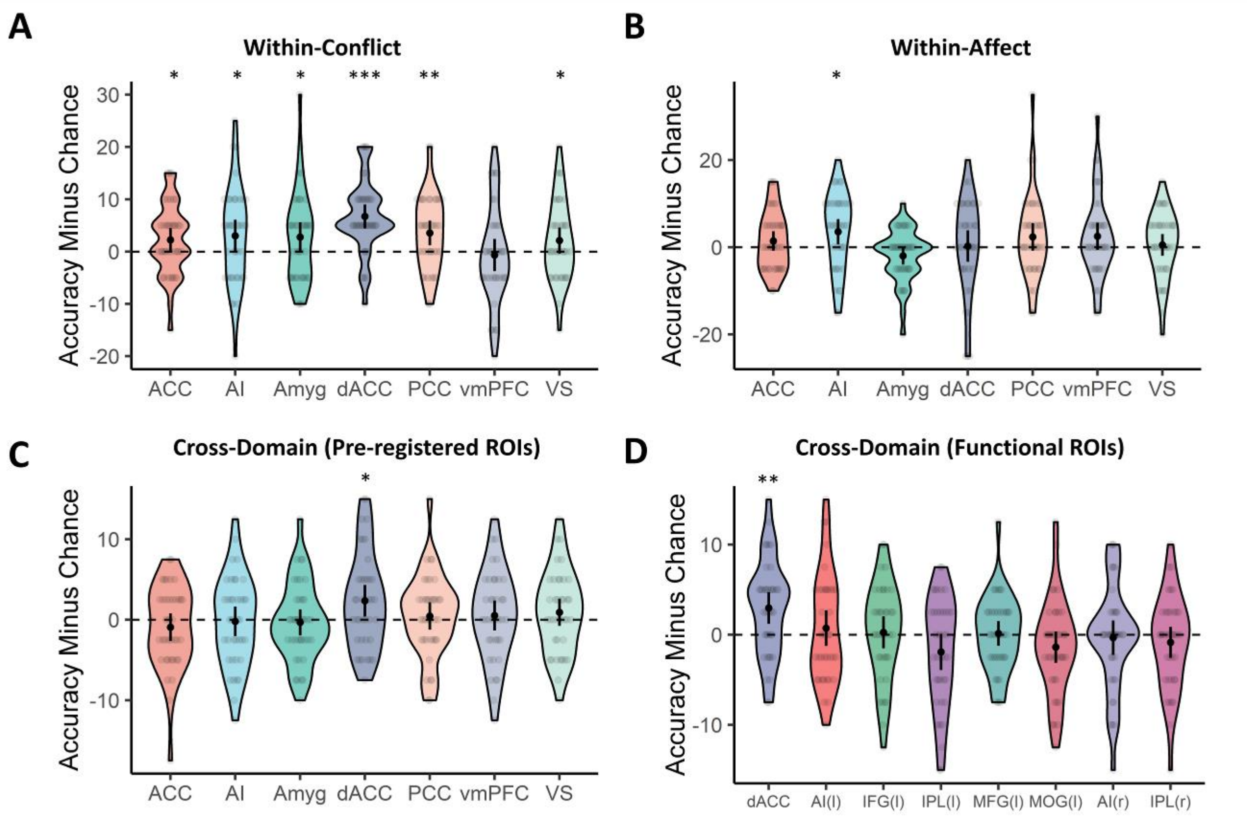

In the "overall decoding" analysis, the classifier was trained and tested in the whole domain regardless of task. **(A)** In the conflict domain, the overall decoding approach replicated above-chance level classification in the ACC (*V*=269, *P*=.024, *BF*=2.07), dACC/pre-SMA (*V*=575, *P*<.001, *BF*>100) and PCC (*V*=575, *P*=.002, *BF*=17.84) in addition to AI (*V*=389, *P*=.024, *BF*=2.02), Amygdala (*V*=374, *P*=.042, *BF*=1.90) and VS (*V*=257, *P*=.047, *BF*=1.24) but not vmPFC (*V*=183, *P*=.068, *BF*=0.13). **(B)** In the affective domain, evidence for above-chance level classification was found in the AI (*V*=407, *P*=.011, *BF*=5.00; all other, *P*>.086, *BF*<0.92). **(C)** The overall cross-domain classification using our pre-registered ROIs again replicated above-chance level classification in the dACC/pre-SMA (*V*=351, *P*=.021, *BF*=4.65; all other, *P*>.319, *BF*<0.22). **(D)** Finally, the overall decoding approach also replicated cross-classification in the dACC/pre-SMA with the functionally defined set of ROIs ($V$=448, $P$=.001, $BF$=41.06; all other $P$>.260, $BF$<0.34). * $P < .05$; ** $P < .01$; *** $P < .001$; black dots and error bars represent mean and ± 95 CI respectively; transparent dots represent individual data points; the shape of the violin shows the distribution of the data. AI(l), left Anterior Insula; IFG(l), left Inferior Frontal Gyrus; IPL(l), left Inferior Parietal Lobule; MFG(l), left Middle Frontal Gyrus; MOG(l), left Middle Occipital Gyrus; AI(r), right Anterior Insula; IPL(r), right Inferior Parietal Lobule.

***Supplementary Figure 2.*** **Within-Affect Decoding**

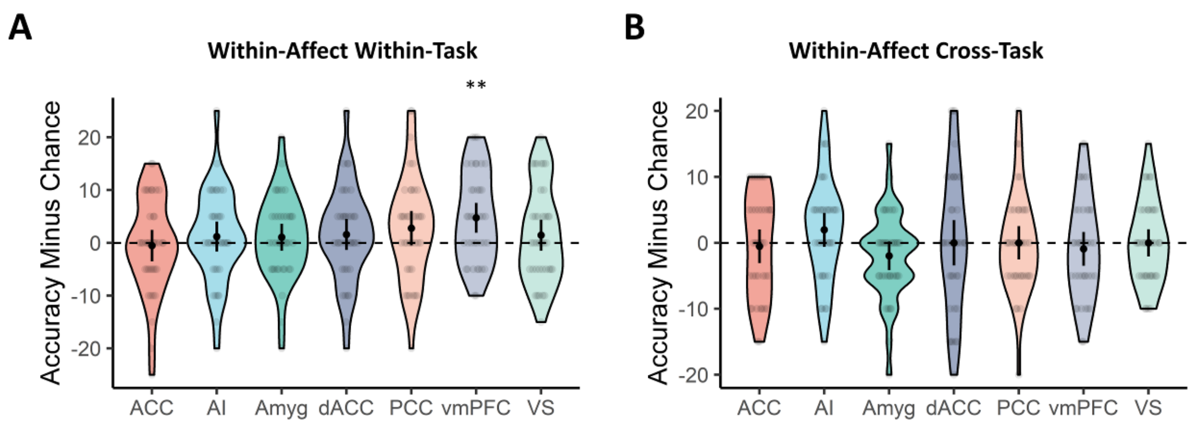

**(A)** Within-task analyses in the affective domain (positive vs. negative) revealed evidence for above chance-level decoding in the vmPFC (*V*=376, *P*=.001, *BF*=39.10; all others, *P*>.073, *BF*<1.26). **(B)** Cross-task analyses did not revealed evidence for successful decoding in any of the ROIs (*P*>.086, *BF*<0.97). The overall decoding of the affect domain can be found in Supplementary Figure 1B and revealed successful decoding in the AI (*V*=407, *P*=.011, *BF*=5.00). * *P* < .05; ** *P* < .01; *** *P* < .001; black dots and error bars represent mean and ± 95 CI respectively; transparent dots represent individual data points; the shape of the violin shows the distribution of the data.

***Supplementary Figure 3.*** **The effect of smoothing on SNR and classification.**

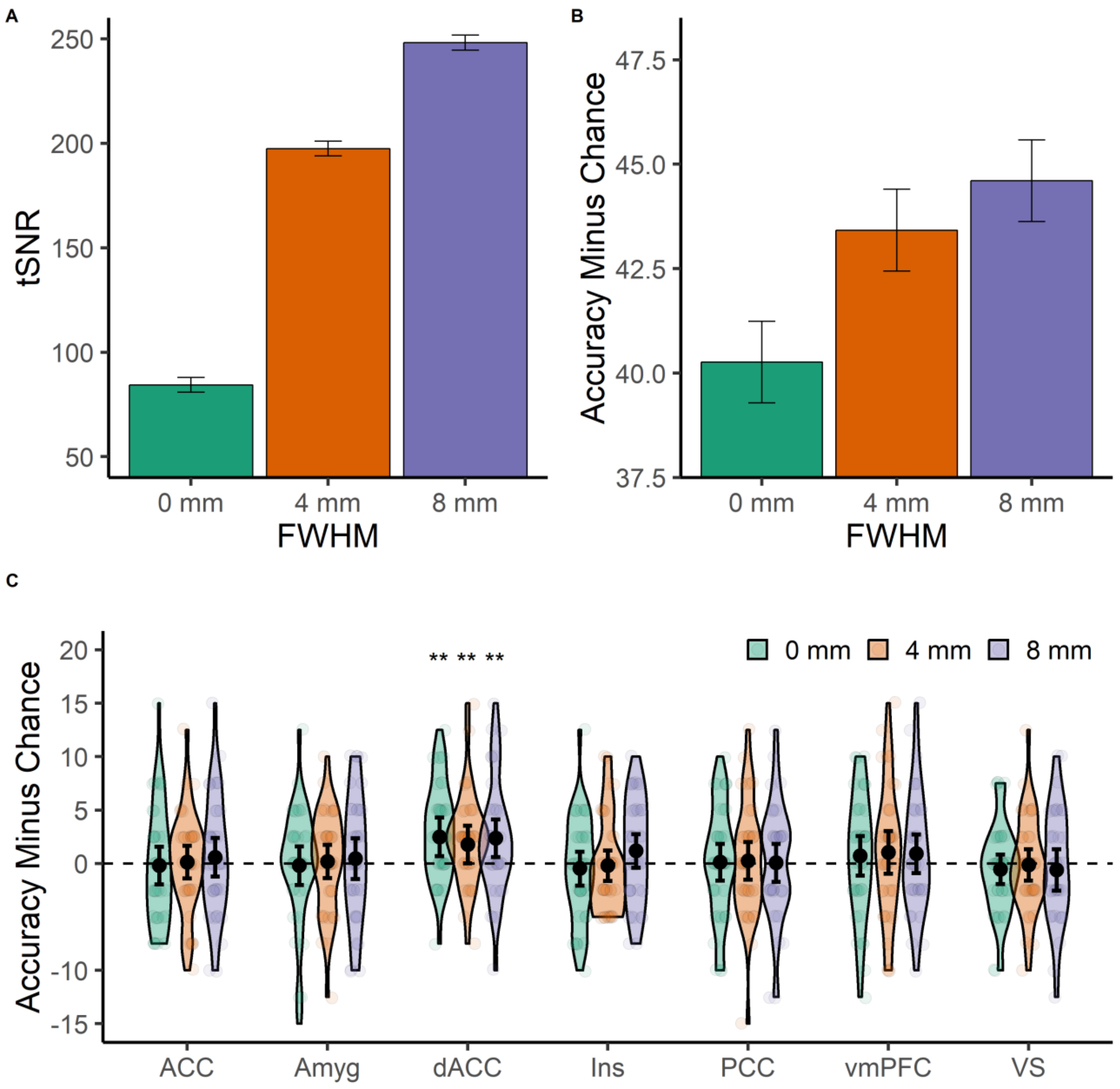

**(A)** Temporal SNR in the primary motor cortex significantly increased with wider smoothing (full width at half-maximum [FWHM], $F_{(1,74)}$=1503, $P$<.001). **(B)** Decoding the laterality of the response hand (left vs right; which was balanced) resulted in higher accuracies with wider smoothing ($F_{(1,74)}$=12.54, $P$<.001). **(C)** Cross-Domain Cross-Task classification decoding accuracy in the main set of ROIs for different smoothing parameters. Changing the smoothing parameters did not chance our main conclusion that dACC/pre-SMA shows above-chance level cross-domain cross-task classification (0mm: $V$=380, $P$=.001, $BF$=9.81, 4mm: $V$=396, $P$=.006, $BF$=2.19, 8mm: $V$=330, $P$=.007, $BF$=8.43). As tSNR and thus reliability increased significantly with smoothing, we chose to report the most reliable data (FWHM 8 mm). * $P < .05$; ** $P < .01$; *** $P < .001$; bars/black dots and error bars represent mean and ± 95 CI respectively; transparent dots represent individual data points; the shape of the violin shows the distribution of the data.

***Supplementary Figure 4.*** **Results using the pre-registered Harvard-Oxford ROIs (with only dACC retrieved from Neurosynth)**

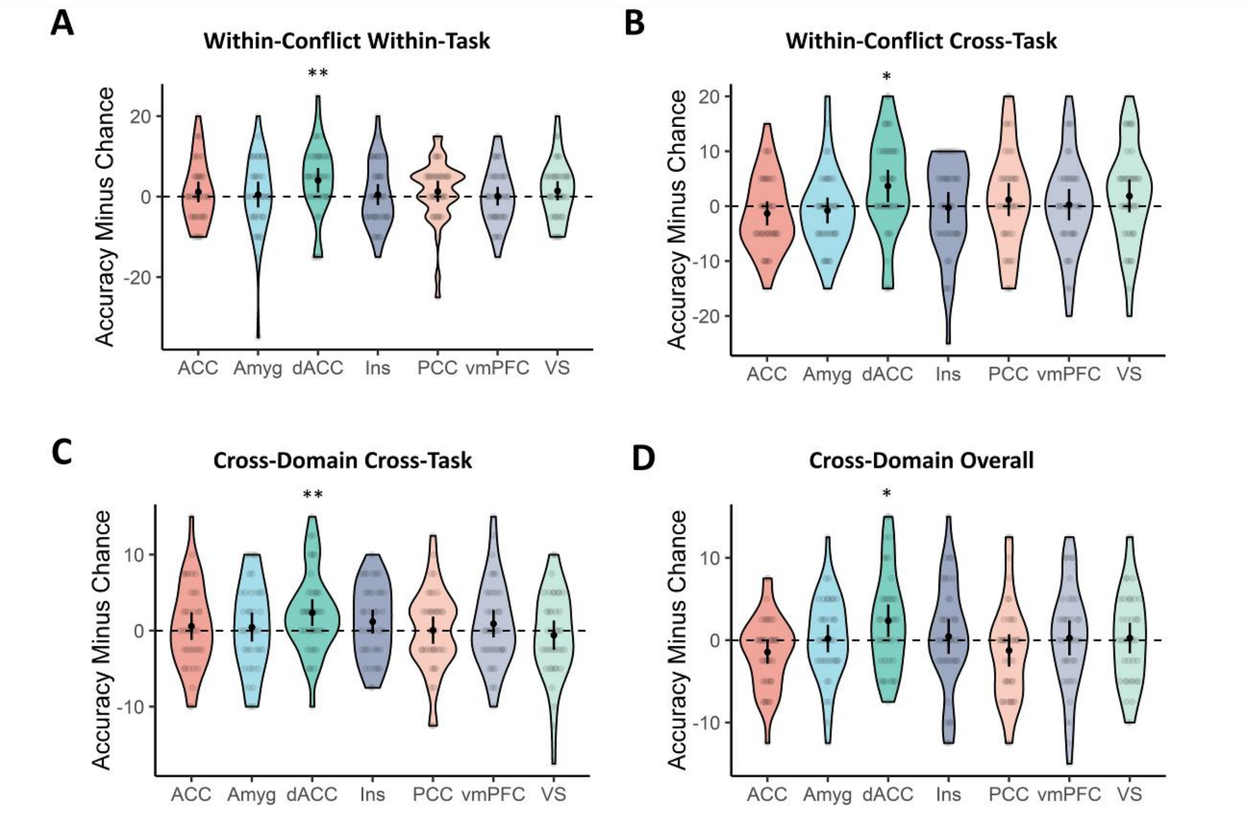

We re-analyzed our results by replacing all non-dACC/pre-SMA NeuroSynth ROIs with our pre-registered Harvard-Oxford atlas ROIs as originally preregistered (ACC, AI, Amyg, PCC, vmPFC, VS). **(A)** The within-task within-conflict decoding revealed only above-chance level classification only in the dACC/pre-SMA ($V$=327, $P$=.008, $BF$=8.48, all other, $P$>.060, $BF$<0.60). **(B)** The same was true for the within-conflict cross-task decoding ($V$=283, $P$=.012, $BF$=5.57, all other, $P$>.117, $BF$<0.64), **(C)** the cross-domain cross-task decoding ($V$=330, $P$=.007, $BF$=8.43, all other, $P$>.066, $BF$<0.94) and **(D)** the overall decoding approach ($V$=351, $P$=.021, $BF$=4.65, all other, $P$>.319, $BF$<0.25) (note: dACC/pre-SMA results are identical to the previously reported results in the main text). * $P$ < .05; ** $P$ < .01; black dots and error bars represent mean and ± 95 CI respectively; transparent dots represent individual data points; the shape of the violin shows the distribution of the data.

***Supplementary Figure 5.*** **Post-scanning Incidental Recognition Memory**

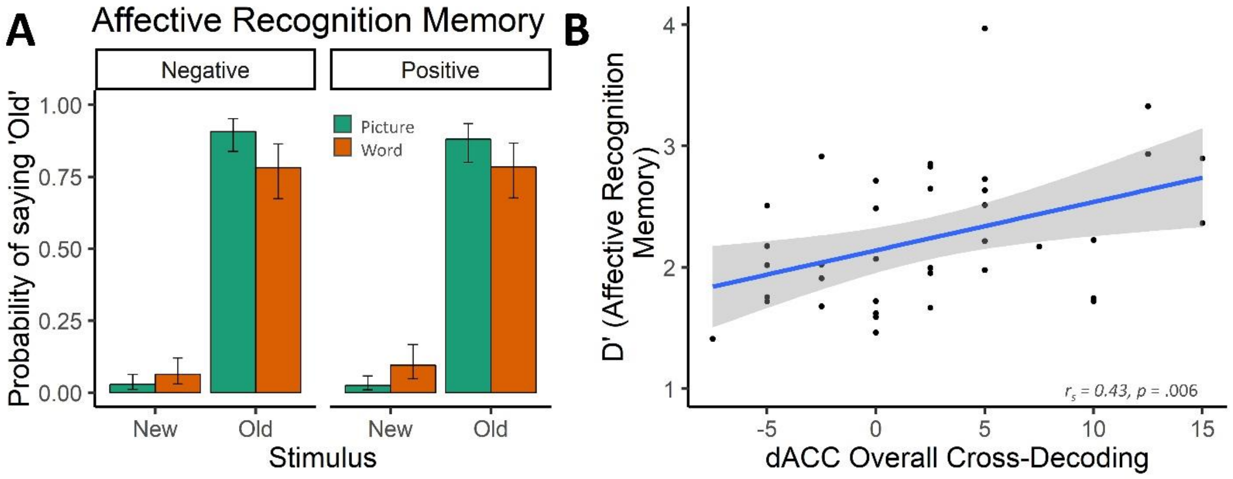

**(A)** After the experiment, subjects performed an unannounced recognition memory test on the affective stimuli. Subjects demonstrated high hit rates and low false alarm rates ($\chi 2$=442, $p$<.001) which was further modulated by affective stimulus type (Pictures > Words, $\chi 2$=16.49, $p$ < .001). Mean ± 95% CI. **(B)** Post-hoc correlation analyses also revealed a significant spearman correlation between the sensitivity index (d-prime) from the post-scanning recognition memory for affective stimuli and overall cross-classification accuracy in the dACC/pre-SMA ROI (*Rs*=0.43, *P*=.006, *BF*=5.02), surviving correction for multiple comparisons (see Methods). This suggests that subjects with better recognition memory were more attentive to the task-irrelevant affective stimuli which made cross-classification more successful. Grey band represents 95% CI.

***Supplementary Figure 6.*** **Searchlight Decoding in the MFC**

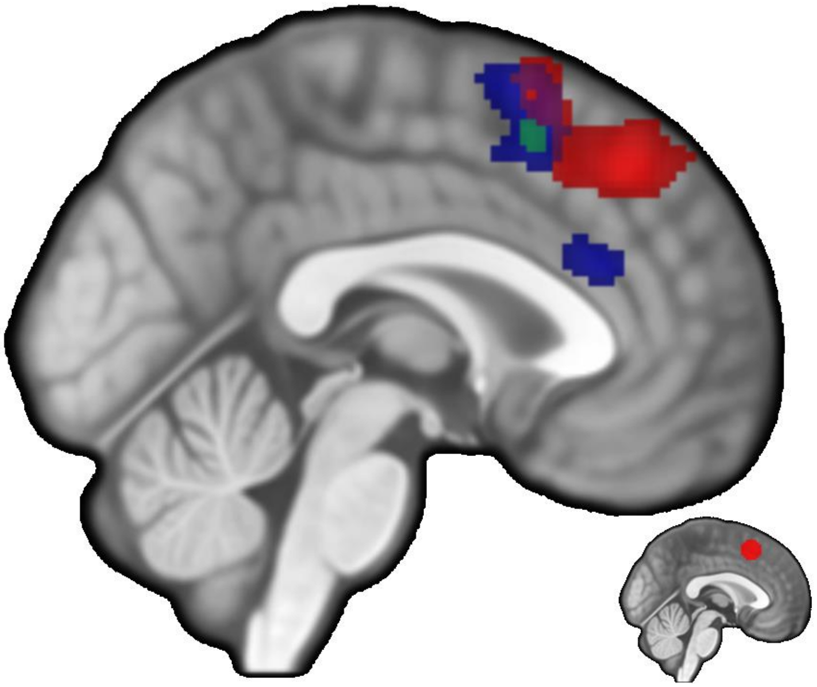

This plot shows the cluster-corrected searchlight decodings restricted to the MFC after a small volume correction (with the same mask as used by Kragel et al., 2018; uncorrected P < .005). Within-affect within-task decoding revealed a large cluster in the dmPFC (N = 865, in red). For the within-conflict within-task searchlight decoding, we show a similar large cluster (N = 3156, in blue). Finally, we found cross-domain cross-task decoding in the MFC, but this cluster (N = 87) did not survive a cluster correction (in green). Note that each of these clusters partially overlap with each other as well as with our main dACC/pre-SMA ROI (bottom-right).

***Supplementary Table 1.*** **Whole-Brain Searchlight Decoding Results.**

| **Anatomical area** | **H** | **X** | **Y** | **Z** | **Voxels** | **Tmax** |
|---|---|---|---|---|---|---|
| *Within-Affect Within-Task Decoding* | | | | | | |
| Left Middle Occipital Gyrus extending into Right Middle Occipital Gyrus | L | -38 | -74 | -4 | 6700 | 10.79 |
| *Within-Affect Cross-Task Decoding* | | | | | | |
| No cluster-corrected activations | | | | | | |
| *Within-Affect Overall Decoding* | | | | | | |
| Middle Occipital Gyrus | L | -32 | -78 | -8 | 2730 | 10.81 |
| Fusiform Gyrus | R | 36 | -42 | -20 | 108 | 7.91 |
| Middle Occipital Gyrus / Middle Temporal Gyrus | R | 46 | -68 | -8 | 602 | 7.72 |
| Insula | R | 38 | 26 | 2 | 208 | 6.63 |
| *Within-Conflict Within-Task Decoding* | | | | | | |
| Inferior Parietal Lobule | L | -28 | -54 | 42 | 467 | 7.36 |
| Middle Frontal Gyrus | L | -44 | 6 | 32 | 779 | 6.88 |
| Middle Occipital Gyrus | L | -26 | -94 | 8 | 226 | 5.99 |
| Pre-SMA | L | -12 | 16 | 54 | 75 | 6.13 |
| *Within-Conflict Cross-Task Decoding* | | | | | | |

| | | | | | | |
|---|---|---|---|---|---|---|
| Superior Parietal Lobule | L | -20 | -62 | 48 | 471 | 8.61 |
| Inferior Frontal Gyrus | L | -34 | 22 | 24 | 42 | 5.96 |
| *Within-Conflict Overall Decoding* | | | | | | |
| Superior Parietal Lobule | L | -26 | -56 | 48 | 2096 | 8.97 |
| Middle Frontal Gyrus extending into pre-SMA | L | -44 | 4 | 38 | 1389 | 8.46 |
| Superior Occipital Gyrus | R | 26 | -70 | 38 | 240 | 6.43 |
| *Cross-Domain Cross-Task Decoding* | | | | | | |
| No cluster-corrected activations | | | | | | |
| *Cross-Domain Overall Decoding* | | | | | | |
| No cluster-corrected activations | | | | | | |

***Supplementary Table 2.*** **Whole-Brain Univariate Results.**

| **Anatomical area** | **H** | **X** | **Y** | **Z** | **Voxels** | **Tmax** |
|---|---|---|---|---|---|---|
| *Conflict Domain Across Both Tasks (incongruent > congruent)* | | | | | | |
| Pre-SMA/dACC | L | -4 | 14 | 50 | 780 | 8.92 |
| Inferior Frontal Gyrus | L | -38 | 22 | 24 | 1040 | 8.38 |
| Superior Parietal Lobule / Precuneus | L | -26 | -58 | 48 | 524 | 8.08 |
| Thalamus | L | -16 | -2 | 16 | 114 | 6.51 |
| Cerebellum | R | 36 | -54 | -30 | 167 | 7.42 |
| *Conflict in the color-word naming task* | | | | | | |
| Middle / Inferior Frontal Gyrus | L | -52 | 10 | 40 | 907 | 8.48 |
| Superior Parietal Lobule / Precuneus | L | -28 | -58 | 50 | 614 | 8.17 |
| Inferior Parietal Lobule | L | -38 | -38 | 42 | 198 | 6.85 |
| Pre-SMA/dACC | L | -6 | 14 | 50 | 190 | 7.51 |
| Middle Frontal Gyrus / Precentral Gyrus | L | -28 | 2 | 68 | 164 | 6.50 |
| *Conflict in the color-circle naming task* | | | | | | |
| Supplementary Motor Area | L | -6 | 4 | 60 | 2 | 5.57 |
| *Affective Domain across tasks (negative > positive)* | | | | | | |
| Left Fusiform Gyrus / Inferior Temporal Gyrus | L | -40 | -44 | -16 | 84 | 7.14 |

| | | | | | | |
|---|---|---|---|---|---|---|
| Middle Occipital Gyrus / Middle Temporal Gyrus | L | -46 | -76 | 2 | 453 | 8.31 |
| Right Inferior Temporal Gyrus | R | 48 | -72 | -2 | 148 | 6.97 |
| *Affect in the color-word naming task* | | | | | | |
| No cluster-corrected activations | | | | | | |
| *Affect in the color-circle naming task* | | | | | | |
| Fusiform Gyrus / Inferior Temporal Gyrus | L | -42 | -44 | -16 | 105 | 8.23 |
| Middle Temporal Gyrus / Middle Occipital Gyrus | R | 50 | -74 | 2 | 244 | 7.76 |
| Middle Temporal Gyrus / Middle Occipital Gyrus | L | -48 | -76 | 8 | 542 | 8.87 |

***Supplementary Table 3.* ANOVA Results for Behavior in the Conflict Domain**

| **Predictor** | **MSE** | ***F*** | **DF** | ***P*** | ***ηG 2*** |
|---|---|---|---|---|---|
| *Reaction Time* | | | | | |
| Intercept | 29712 | 2661.37 | 1, 37 | < .001 | .98 |
| Condition | 1628 | 149.81 | 1, 37 | < .001 | .16 |
| Task | 3623 | 2.71 | 1, 37 | .110 | .007 |
| Condition * Task | 885 | 35.55 | 1, 37 | < .001 | .02 |
| *Accuracy* | | | | | |
| Intercept | .01 | 27065 | 1, 37 | < .001 | >.99 |
| Condition | <.01 | 11.72 | 1, 37 | .002 | .05 |
| Task | <.01 | 0.02 | 1, 37 | .898 | < .001 |
| Condition * Task | <.01 | 0.30 | 1, 37 | .586 | .001 |

*Note*. ηG 2: Generalized Eta Squared; MSE: Mean Square Error.

***Supplementary Table 4.*** **ANOVA Results for Behavior in the Affect Domain**

| **Predictor** | **MSE** | ***F*** | **DF** | ***P*** | ***ηG 2*** |
|---|---|---|---|---|---|
| *Reaction Time* | | | | | |
| Intercept | 30484 | 2557.10 | 1, 37 | < .001 | .98 |
| Condition | 290 | 5.47 | 1, 37 | .025 | .001 |
| Task | 2060 | 3.61 | 1, 37 | .065 | .006 |
| Condition * Task | 229 | 4.27 | 1, 37 | .046 | < .001 |
| *Accuracy* | | | | | |
| Intercept | .01 | 24846.85 | 1, 37 | < .001 | > .99 |
| Condition | <.01 | 0.09 | 1, 37 | .765 | < .001 |
| Task | <.01 | 1.85 | 1, 37 | .181 | .005 |
| Condition * Task | <.01 | 3.12 | 1, 37 | .085 | .005 |

*Note*. ηG 2: Generalized Eta Squared; MSE: Mean Square Error.

***Supplementary Table 5.*** **Correlations Between Individual Differences Measures and Cross-domain Cross-task Decoding in dACC/pre-SMA**

| **Measure** | ***R*** | ***P*** |
|---|---|---|
| Positive Affect (PANAS) | -0.12 | .73 |
| Negative Affect (PANAS) | 0.06 | .46 |
| Need For Cognition (NFC) | 0.04 | .81 |
| Attention Impulsivity (BIS11) | -0.03 | .83 |
| Motor Impulsivity (BIS11) | 0.01 | .93 |
| Non-planning (BIS11) | -0.19 | .26 |
| BAS Drive | -0.03 | .85 |
| BAS Fun | 0.04 | .81 |
| BIS | 0.23 | .17 |

***Supplementary Table 6.*** **Correlations Between Individual Differences Measures and Cross-domain Cross-task Decoding in vmPFC**

| **Measure** | ***R*** | ***P*** |
|---|---|---|
| Positive Affect (PANAS) | 0.02 | .89 |
| Negative Affect (PANAS) | -0.02 | .93 |
| Need For Cognition (NFC) | -0.12 | .48 |
| Attention Impulsivity (BIS11) | 0.04 | .81 |
| Motor Impulsivity (BIS11) | -0.17 | .31 |
| Non-planning (BIS11) | -0.04 | .80 |
| BAS Drive | -0.12 | .47 |
| BAS Fun | -0.16 | .34 |
| BIS | 0 | .99 |
